# pWGBSSimla: a profile-based whole-genome bisulphite sequencing data simulator incorporating methylation QTLs, allele-specific methylations and differentially methylated regions

**DOI:** 10.1101/390633

**Authors:** Ren-Hua Chung, Chen-Yu Kang

## Abstract

**Motivation:** DNA methylation plays an important role in regulating gene expression. There has been growing interest in investigating the roles that genetic variants play in changing the methylation levels (i.e., methylation quantitative trait loci or meQTLs), how methylation regulates the imprinting of gene expression (i.e., allele-specific methylation or ASM), and the differentially methylated regions (DMRs) among different cell types. However, none of the current simulation tools can generate whole-genome bisulphite sequencing (WGBS) data while modeling meQTLs, ASM, and DMRs.

**Results:** We developed pWGBSSimla, a profile-based WGBS data simulator, which simulates WGBS data for 29 cell types based on real data. meQTLs and ASM are modeled based on the block structures of methylation status at CpGs, and DMRs are simulated based on observations of methylation rates in real data.

**Availability:** pWGBSSimla is available at http://omicssimla.sourceforge.io.

## Introduction

DNA methylation plays an important role in cellular development and disease. Methylation quantitative trait loci (meQTLs), which are genetic variants (e.g., SNPs) affecting methylation levels, are associated with changes in chromatin and gene expression and with disease risk (Banovich, et al., 2014). Moreover, allele-specific methylation (ASM), which paternal and maternal alleles have different probabilities of being methylated, can regulate imprinting of gene expression. Furthermore, differentially methylated regions (DMRs), where differences in the methylation levels are observed across various cell and tissue types, are involved in tissue- or cell-specific gene expression.

The current simulators for whole-genome bisulphite sequencing (WGBS) data, such as DNemulator (Frith, et al., 2012) and WGBSSuite (Rackham, et al., 2015), focus on simulating the methylation levels of CpGs, and meQTLs, ASM, and DMRs are not modeled. Here, we developed a profile-based WGBS data simulator (pWGBSSimla), which simulates WGBS data based on the profiles for 29 human cell types, such as blood CD4 and brain cells. Based on the observations that methylated and unmethylated CpGs are clustered in the human genome (Su, et al., 2013), methylated blocks (MBs), unmethylated blocks (UBs), and fuzzily methylated blocks (FBs) were defined in the profiles. By integrating the DNA sequence simulator SeqSIMLA2 (Chung, et al., 2015) with pWGBSSimla, SNP and WGBS data are both simulated for unrelated/related cases and controls, and the meQTLs affecting CpGs in MBs, UBs, or FBs are modeled. Moreover, because an FB may result from a local ASM (Shoemaker, et al., 2010), correlated ASM levels are simulated in the same FB. Finally, DMRs are generated by simulating the same methylation region for different cell types.

## Methods

### Profile generation

We used the 41 WGBS datasets for 29 human cell types (Ziller, et al., 2013) for the profile generation. Several QC steps were performed for the methylation calls. CpGs remained across all 41 samples after QC were used. For each cell type, the methylation rate for each CpG and the distances between adjacent CpGs were calculated. Then the hotspot extension algorithm (Su, et al., 2013) was used to identify MBs, UBs, and FBs. Finally, the read depths were classified for each block type. More details about the profile generation can be found in the Supplementary Materials.

### Simulation of WGBS data

Given a user-specified genomic region and cell type, CpGs and the distances between them were first generated based on the profiles. The read depth of a CpG in an MB, UB, or FB in a sample was then randomly sampled from the read depths of the corresponding block type in the profile. Finally, the methylated read count of CpG *i* in a sample is simulated based on a Binomial distribution with parameters (*n*_*i*_, *p*_*i*_), where *n*_*i*_ is the read depth and *p*_*i*_ is the methylation rate of CpG *i* from the profile.

### Simulation of meQTLs, ASMs, and DMRs

It was observed that a meQTL can affect the methylation levels at multiple CpGs in a local region up to 3 kb (Banovich, et al., 2014). Moreover, it has been shown that the ASM at adjacent CpGs can be highly correlated (Shoemaker, et al., 2010). Therefore, given a user-specified meQTL at a SNP, an MB, UB, or FB is randomly sampled to be influenced by the meQTL. The methylated read counts for CpGs located in up to 3 kb of the block are simulated based on the same genotype-specific methylation probabilities, which are *p*_*i*_, *p*_*i*_(1+*f*_*i*_), and *p*_*i*_(1+2*f*_*i*_) for genotypes *AA*, *Aa*, and *aa*, respectively, where *A* is the reference allele at meQTL *i*. The simulation of SNP genotypes is described in the next section. More details for determining *f*_*i*_ can be found in the Supplementary Materials. Furthermore, given the proportion of ASM specified by the user, the FBs are randomly sampled to have ASM. The methylated read counts for paternal and maternal alleles are determined by *p*_*i*_ × *r^father^* and *p^i^* × *r^mother^*, respectively, where *r^father^* and *r^mother^* are the ratios of the father- and mother-specific methylation rates, respectively, relative to *p^i^*. Note that ***p*** for CpGs in the same block is determined using a multivariate beta distribution to model the correlation of ASM. More details can be found in the Supplementary Materials. Finally, because the profiles for 29 different cell and tissue types are compiled, it is straightforward to generate DMRs by simulating the same genomic region using profiles for two or more cell/tissue types.

### Simulation of methylation difference in cases and controls

The SeqSIMLA2 algorithm is first used to simulate SNP genotypes in affected and unaffected individuals. A brief review of SeqSIMLA2 is provided in the Supplementary Materials. Some of the SNPs are specified as meQTLs by the user. Given a user-specified proportion of methylation difference for a block type (*d^t^*, where *t* is MB, UB or FB), blocks with type *t* are randomly sampled, so the total number of CpG sites in the sampled blocks is approximately *d^t^* × total CpG sites, similar to the algorithm in WGBSSuite. The methylation probability for CpG *i* is (*p*_*i*_ + θ) in half of the sampled blocks and is (*p*_*i*_ − θ) in the other half of the sampled blocks for affected individuals, where *θ* is the difference in the methylation rate between the cases and controls.

## Results

Figure 1 shows the percentages of CpGs in MBs, UBs, and FBs and the distributions of block sizes (i.e., numbers of CpGs in blocks) for the liver cell. Consistent with previous observations (Su, et al., 2013), most of the sites (>97%) are in MBs and UBs. There are small intersections of CpGs between the block types. The UBs have the largest median of sizes (88), followed by MBs (80) and FBs (17). Figure 2A shows the methylation rates at a meQTL. As expected, differences in the methylation rates both between the genotypes and between affected and unaffected individuals were observed. Figure 2B shows pairwise correlations of methylation levels in an ASM region. As seen in the Figure, a local “methylation LD block”, as previously observed (Shoemaker, et al., 2010), can be simulated. Figure 2C shows the difference in the mean methylation levels between the liver and colon cells in a DMR (the OCT4 locus) that was previously reported (Ziller, et al., 2013). Similar to their observations, there is a small difference in the mean methylation levels in CpG islands and CTCF-binding sites, but larger differences are observed in promoters and within genes. The results for other cell types for Figure 1 and more details for the simulations for Figure 2 are provided in the Supplementary Materials. In conclusion, our results demonstrate that pWGBSSimla simulates WGBS data while properly modeling meQTLs, ASM, and DMRs.

**Fig. 1.**
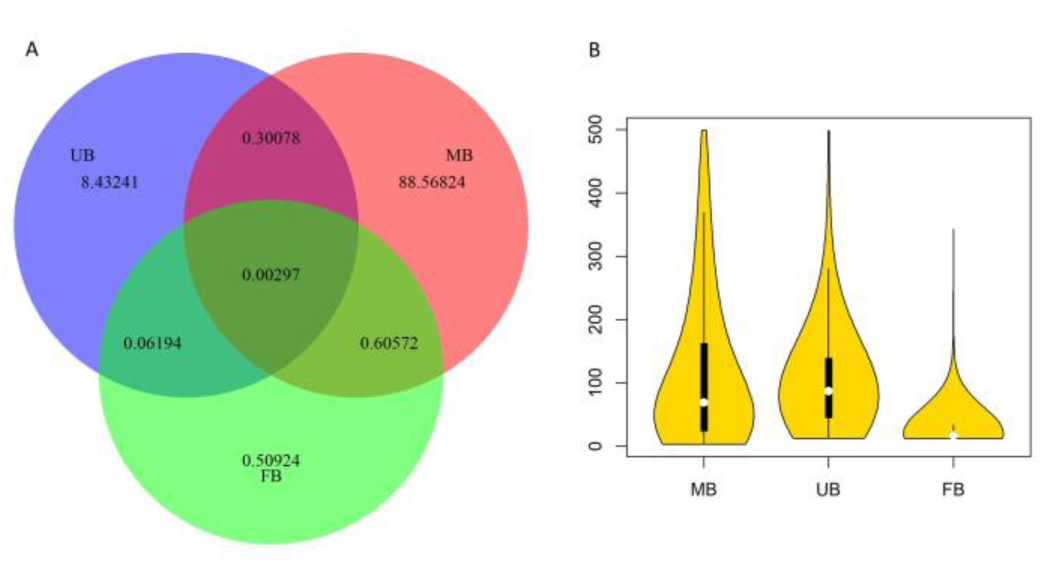
(A) The percentages of CpGs covered by UBs, MBs, and FBs. (B) Violin plot for the distribution of block size for each block type.

**Fig. 2.**
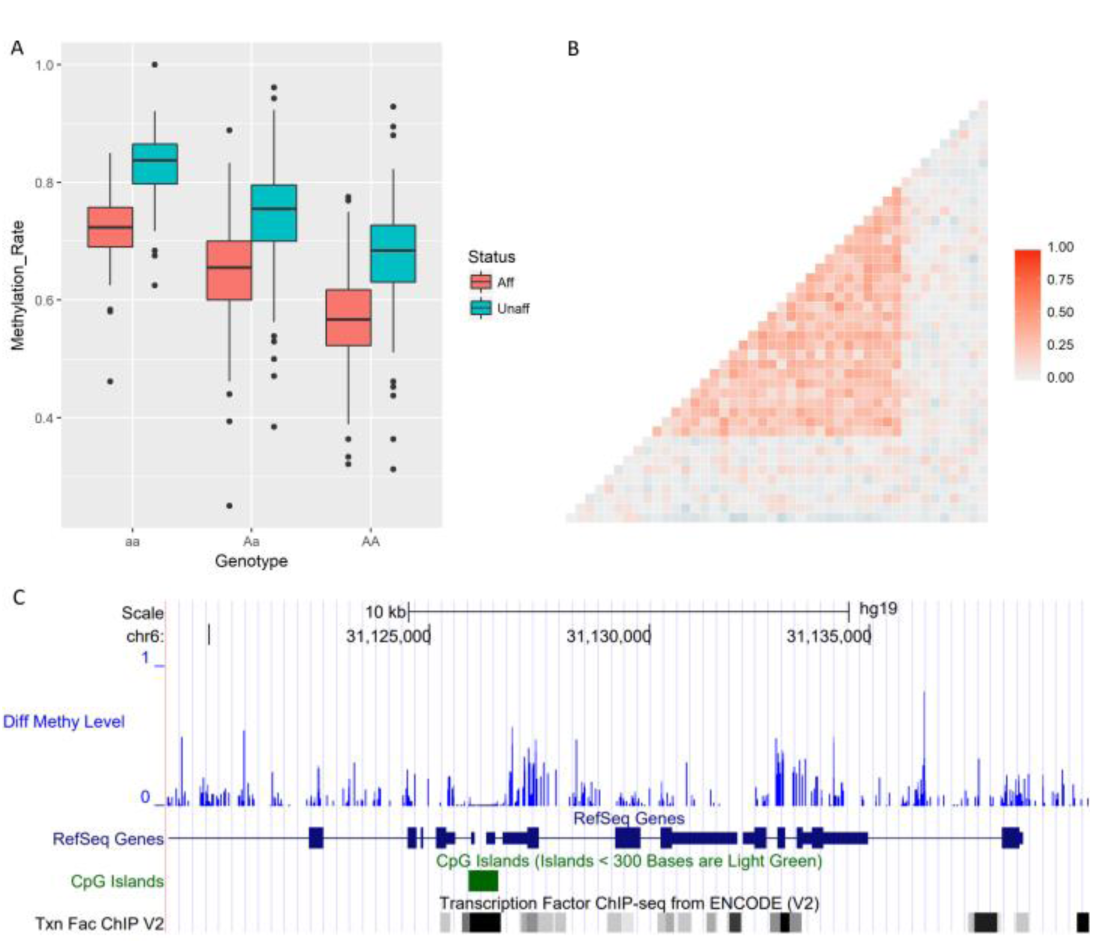
(A) Distributions of methylation rates at a meQTL. (B) Pairwise correlations of methylation levels in an ASM region. (C) Difference in methylation levels between the liver and colon tissues at the OCT4 locus (chr6: 31,119,000 - 31,140,000).

## Acknowledgments

This work has been supported by the grants from the Ministry of Science and Technology (MOST 106-2221-E-400-005-MY3) in Taiwan.

